# MICOS and F_1_F_O_-ATP synthase crosstalk is a fundamental property of mitochondrial cristae

**DOI:** 10.1101/2021.04.01.438160

**Authors:** Lawrence Rudy Cadena, Ondřej Gahura, Brian Panicucci, Alena Zíková, Hassan Hashimi

## Abstract

Mitochondrial cristae are polymorphic invaginations of the inner membrane that are the fabric of cellular respiration. Both the Mitochondrial Contact Site and Cristae Organization System (MICOS) and the F_O_F_1_-ATP synthase are vital for sculpting cristae by opposing membrane bending forces. While MICOS promotes negative curvature at cristae junctions, dimeric F_O_F_1_- ATP synthase is crucial for positive curvature at cristae rims. Crosstalk between these two complexes has been observed in baker’s yeast, the model organism of the Opisthokonta supergroup. Here, we report that this property is conserved in *Trypanosoma brucei*, a member of the Discoba supergroup that separated from Opisthokonta ∼2 billion years ago. Specifically, one of the paralogs of the core MICOS subunit Mic10 interacts with dimeric F_1_F_O_-ATP synthase, whereas the other core Mic60 subunit has a counteractive effect on F_1_F_O_- ATP synthase oligomerization. This is evocative of the nature of MICOS-F_1_F_O_-ATP synthase crosstalk in yeast, which is remarkable given the diversification these two complexes have undergone during almost 2 eons of independent evolution. Furthermore, we identified a highly diverged trypanosome homolog of subunit e, which is essential for the stability of F_1_F_O_-ATP synthase dimers in yeast. Just like subunit e, it is preferentially associated with dimers, interacts with Mic10 and its silencing results in severe defects to cristae and disintegration of F_1_F_O_-ATP synthase dimers. Our findings indicate that crosstalk between MICOS and dimeric F_1_F_O_-ATP synthase is a fundamental property impacting cristae shape throughout eukaryotes.

**Importance:** Mitochondria have undergone profound diversification in separate lineages that have radiated since the last common ancestor of eukaryotes some eons ago. Most eukaryotes are unicellular protists, including etiological agents of infectious diseases like *Trypanosoma brucei*. Thus, the study of a broad range of protists can reveal fundamental features shared by all eukaryotes and lineage-specific innovations. Here we report that two different protein complexes, MICOS and F_1_F_O_-ATP synthase, known to affect mitochondrial architecture, undergo crosstalk in *T. brucei*, just as in baker’s yeast. This is remarkable considering that these complexes have otherwise undergone many changes during their almost two billion years of independent evolution. Thus, this crosstalk is a fundamental property needed to maintain proper mitochondrial structure even if the constituent players considerably diverged.

## Introduction

Mitochondria are ubiquitous organelles that play a central role in cellular respiration in aerobic eukaryotes alongside other essential processes, some of which are retained in anaerobes (1). These organelles have a complex internal organization. While the mitochondrial outer membrane is smooth, the inner membrane is markedly folded into invaginations called cristae, the morphological hallmark of the organelle in facultative and obligate aerobes (2, 3). Cristae are enriched with respiratory chain complexes that perform oxidative phosphorylation (4, 5), and eukaryotes that cannot form these complexes lack these ultrastructures (1, 6).

Mitochondrial morphology differs between species, tissues, and metabolic states. Nevertheless, these variations are derived from common structural features, some of which were likely inherited from endosymbiotic α-proteobacterium that gave rise to the organelle (1, 7,8). Cristae contribute to this variety as they can assume different shapes, such as plate-like lamellar cristae, which decorate yeast and animal mitochondria, and the paddle-like discoidal cristae seen in discoban protists (9). These shapes are at least in part determined by protein complexes embedded within the cristae membranes.

The Mitochondrial Contact Site and Cristae Organization System (MICOS) is a hetero-oligomeric protein complex that is responsible for the formation and maintenance of cristae junctions, narrow attachment points of cristae to the rest of the inner membrane (10, 11). Cristae junctions serve as diffusion barriers into and out of cristae, as these structures cease to act as autonomous bioenergetic units upon MICOS ablation (12). The two core MICOS subunits Mic10 and Mic60, which are both well conserved throughout eukaryotes (13, 14), have demonstrated membrane modeling activity that contributes to constriction at cristae junctions (15-18).

The F_1_F_O_-ATP synthase (henceforth ‘ATP synthase’) dimers also influence cristae shape (19). Dimeric ATP synthase assemble into rows or other oligomeric configurations that promote positive curvature at crista rims (20, 21). In yeast and animals, both belonging to the eukaryotic supergroup Opisthokonta (22), ATP synthase dimerization depends on membrane- embedded F_O_ moiety subunits e and g (23-25). Although these subunits do not directly contribute to intermonomer contacts, deletion of either of them hinders dimer formation (20) and subsequently results in the emergence of defective cristae (25, 26).

Crosstalk between the crista shaping factors MICOS and ATP synthase has been demonstrated in yeast. A fraction of Mic10 physically interacts with ATP synthase dimers, presumably via subunit e, and overexpression of Mic10 leads to stabilization of ATP synthase oligomers (27, 28). Even more pronounced accumulation of ATP synthase oligomers was observed upon deletion of the Mic60 gene, but no physical interaction has been reported between Mic60 and any ATP synthase subunits (29). This interplay between MICOS and ATP synthase is a remarkable and still unexplained phenomenon, as both complexes play critical, yet apparently antithetical roles in shaping the inner membrane.

Here, we ask whether crosstalk between MICOS and ATP synthase dimers is a fundamental property of cristae. We employed the protist *Trypanosoma brucei*, a model organism that is part of supergroup Discoba, which diverged from Opisthokonta ∼1.8 billion years ago (22, 30). Because of their extended independent evolution, discoban MICOS (30-32) and ATP synthase (33-36) differ radically from their opisthokont counterparts. Discoban MICOS has two Mic10 paralogs and an unconventional Mic60, which lacks the C-terminal mitofilin domain, characteristic for other Mic60 orthologs (13-14). Furthermore, unlike opisthokont MICOS, which is organized into two integral membrane sub-complexes, each assembled around a single core subunit, trypanosome MICOS is composed of one integral and one peripheral sub-complex. Discoban ATP synthase exhibits type IV dimer architecture, different from the canonical type I dimers found in opisthokonts (20-33), and has a dissimilar F_O_ moiety, which lacks obvious homologs of subunit e or g (37).

## Results

### Depletion of trypanosome Mic60 affects oligomerization of F_1_F_O_-ATP synthase independent of other MICOS proteins

In *T. brucei*, ablation of conserved MICOS components, Mic60 or both Mic10 paralogs simultaneously, resulted in impaired submitochondrial morphology characterized by elongated cristae adopting arc-like structures (31). Because in budding yeast, the knockout of Mic60 affects oligomerization of ATP synthase (26), a major contributor to cristae organization, we asked whether ATP synthase plays a role in the changes in mitochondrial ultrastructure observed after Mic10 and Mic60 depletion. After inducible RNAi silencing of Mic60 (Mic60↓), we observed increased levels of ATP synthase subunits β, a component of the catalytic F_1_ sector, oligomycin sensitivity conferral protein (OSCP), a subunit of the peripheral stalk, and ATPTb2, a distant homolog of opisthokont subunit d. Simultaneously, levels of Mic10-1 were mildly reduced (Fig. 1A). Notably, RNAi induction of Mic10-2 in the Mic10-1 knock-out cell line (*ΔMic10-1*:Mic10-2↓) resulted in the same cristae defects as in Mic60↓ cells (30), but not in detectable changes in steady state levels of ATP synthase subunits (Fig. S1A). Thus, the accumulation of ATP synthase subunits documented upon Mic60 knock-down cannot be explained as a general consequence of morphologically elongated and detached cristae.

**Figure 1.**
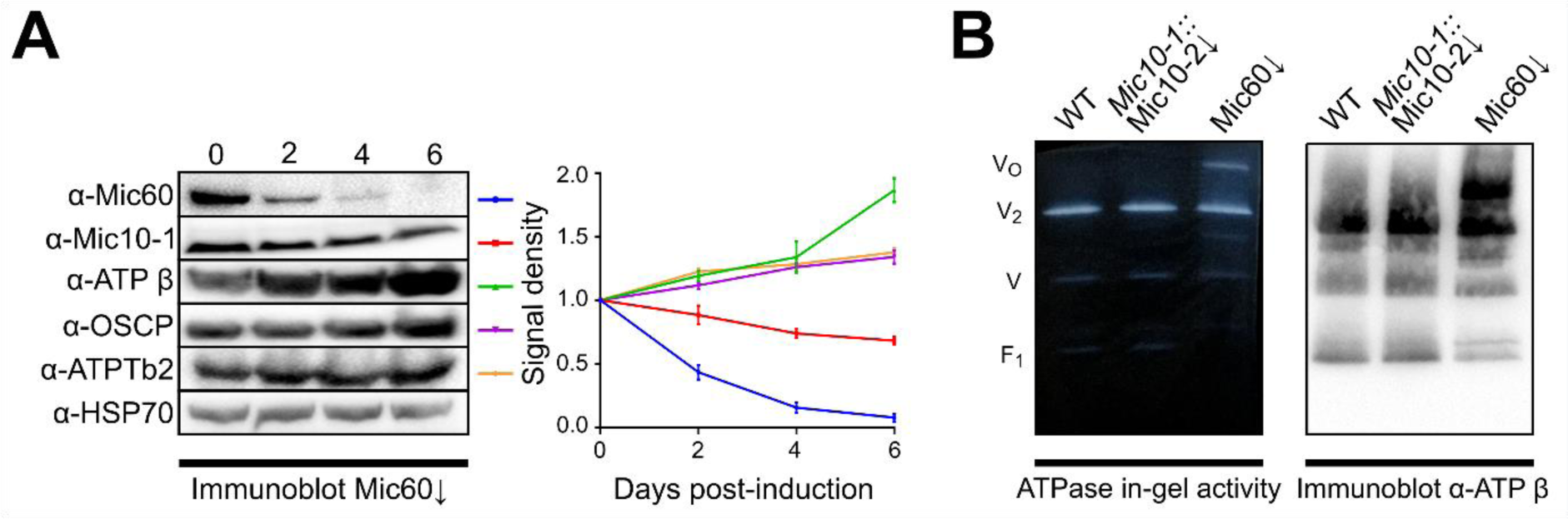
F_O_F_1_ -ATP synthase oligomers are stabilized upon Mic60 depletion. A) Representative immunoblots of whole cell lysate from Mic60↓ cells over a 6-day course of RNAi induction with antibodies indicated on left and densiometric quantification of immunoblots done in triplicate, with signal normalized to unaffected HSP70 (error bars, SD). Days post-induction are shown above immunoblots. Line colors to the right of each immunoblot label antibody signal intensities shown in the graph. Signal density is plotted in arbitrary units B) Blue native polyacrylamide using 1.5% DDM-solubilized mitochondria from *ΔMic10-1*:Mic10-2↓ and Mic60↓ RNAi cells at 5 days post-induction compared to wild type (WT). ATP hydrolysis in-gel activity staining shown on the left and an immunoblot probed with α-ATP synthase β-subunit antibody on the right. V_O_, oligomer; V_2_, dimer; V, monomer; F_1_, free F_1_ moiety.

The increased abundance of ATP synthase subunits prompted us to investigate if the depletion of Mic60 affects the oligomeric state of ATP synthase. RNAi induction in Mic60↓, but not in *ΔMic10-1*:Mic10-2↓ cells, led to the stabilization of ATP synthase oligomers compared to wild type cells, as documented by immunodetection of ATP synthase complexes in 1.5% n- Dodecyl-β-D-Maltoside (DDM) solubilized mitochondria resolved by blue native polyacrylamide gel electrophoresis (BN-PAGE; Fig. 1B). ATP hydrolysis in-gel activity staining demonstrated that the oligomers are enzymatically active. To confirm that the accumulation of ATP synthase dimers was exclusive to Mic60↓, we screened the effect of depletion of two kinetoplastid specific subunits of MICOS known to disrupt cristae, Mic32 and Mic20. Upon their depletion, no changes in ATP synthase oligomerization were observed on native gels (Fig. S1B). In conclusion, we present evidence that Mic60 depletion in *T. brucei* alters the oligomeric state of ATP synthase independently of the defects to cristae.

### Trypanosome Mic10-1 interacts with F_1_F_O_-ATP synthase

To analyze whether any of the Mic10 paralogs or Mic60 interact with ATP synthase, the F_O_ moiety ATPTb2 was C-terminally V5-epitope-tagged and used for co-immunoprecipitation (co-IP) from chemically crosslinked mitochondrial lysates. To facilitate immunocapture proteins interacting with ATPTb2-V5, hypotonically isolated mitochondria were crosslinked with dithiobis(succinimidyl propionate) (DSP) prior to solubilization. These were subsequently incubated with mouse α-V5 antibody conjugated to protein G Dynabeads to immunoprecipitate (IP) the tag. After extensive washing, eluted proteins that co-IP with ATPTb2 were separated via protein electrophoresis, blotted and the co-IP eluate was probed for the presence of Mic10- 1 (molecular weight of this and other proteins investigated here are given in table S1). Mic10- 1 was shown to co-IP with ATPTb2, yielding two discernible bands at approximately 35 kDa and 70 kDa (Fig. 2A). While uncrosslinked Mic10-1 is prominent in the input, it is completely absent in the eluate, suggesting that the protein can co-immunoprecipitate with ATPTb2 only when permanently linked to an unknown partner. We speculate that the ∼35 kDa band corresponds to an adduct between Mic10-1 and the unknown partner, and the ∼70 kDa band may represent the same adduct crosslinked to ATPTb2.

**Figure 2.**
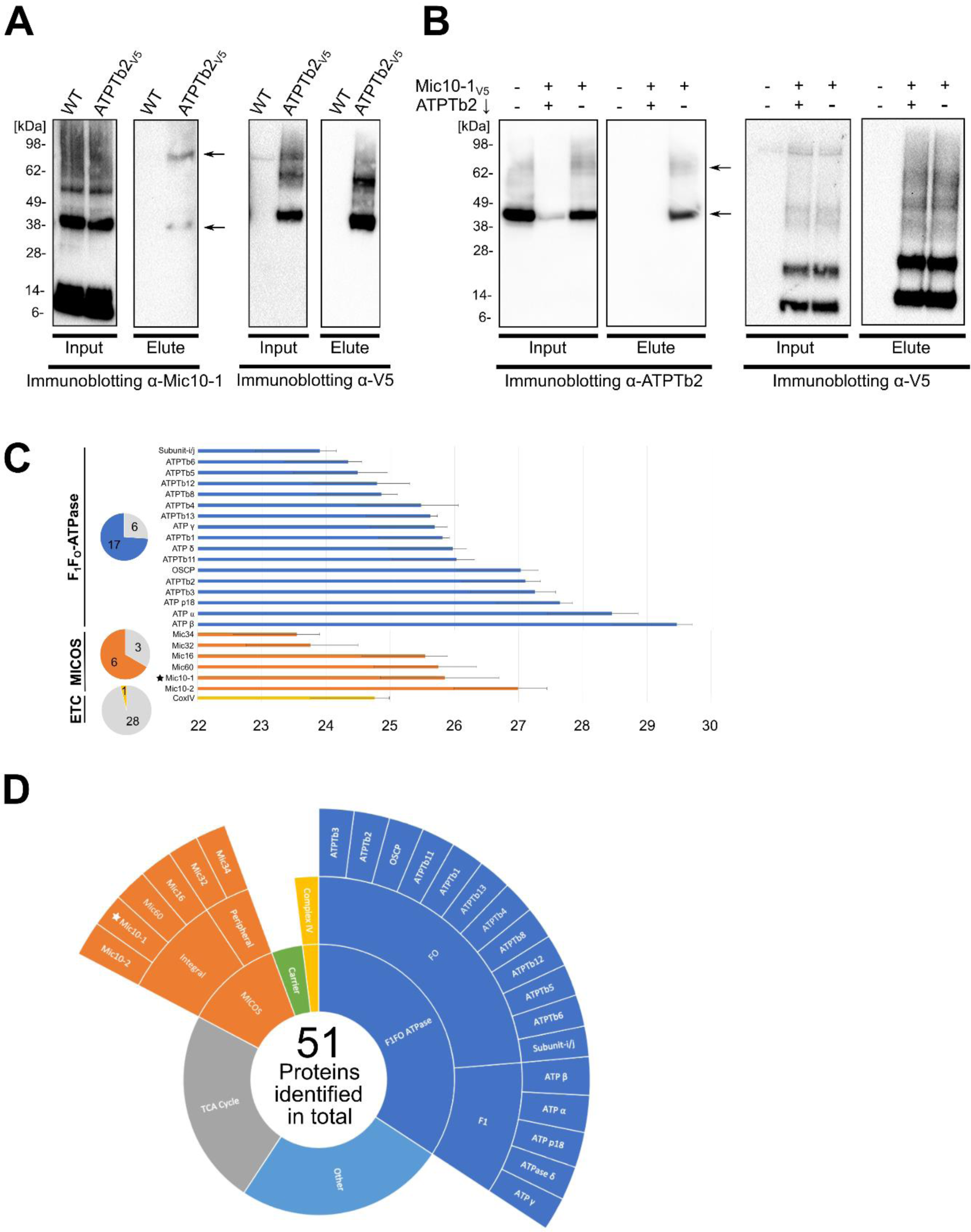
Mic10-1 interacts with F_O_F_1_-ATP synthase. A) Immunoprecipitation (IP) of wild type (WT) and ATPTb2-V5 mitochondria crosslinked with 80 μM DSP with anti-V5 antibody. Input (10% of elute) and eluates were resolved by SDS-PAGE and immunoblotted with antibody against Mic10-1 and V5-peptide. Arrows indicate discernible bands corresponding to ∼35 kDa and ∼70 kDa. B) IP of WT, Mic10-1-V5:ATPTb2↓ and Mic10-1-V5 mitochondria crosslinked with 80 μM DSP. Input (10% of elute) and eluates were resolved by SDS-PAGE and immunoblotted against α-ATPTb2 and V5-peptide. Arrows indicate discernible bands corresponding to ∼40 kDa and ∼70 kDa. C) Summary of oxidative phosphorylation and MICOS complexes subunits that co-IP with Mic10-1-V5 identified by mass spectroscopy. Proteins in dark blue belong to the ATP synthase complex, orange to the MICOS complex, and yellow to the electron transport chain (ETC). The naming for the ATP synthase subunits was taken from Perez et al., 2014 and Gahura et al., 2021 (45, 37). Star indicates bait protein Mic10-1- V5. Intensity score, x axis. (n = 3; error bars, SD). Pie charts on the right depict the portion of proteins identified vs total number of subunits in each respective complex. Other proteins fitting the criteria are given in Fig. S2B. D) Sunburst schematic of all proteins that co-IP with Mic10-1-V5 identified above the established threshold and criteria described in the main text. Subunits of the ATP synthase complex are shown in dark blue and are further distinguished into the F_1_ and F_O_ moieties. Proteins belonging to the MICOS complex are shown in orange and are further separated into integral or peripheral membrane proteins. ETC complex cytochrome c oxidase is shown in yellow. Mitochondrial carriers in green and components of the tricarboxylic acid cycle (TCA) in grey. Other proteins which could not be categorized into a single group are depicted as ‘Other’.

In order to verify the interaction between Mic10-1 and ATPTb2, we performed a reciprocal pulldown of C-terminal V5-epitope-tagged Mic10-1. Indeed, we were able to capture ATPTb2 in the eluted protein fraction with bands present at the 40 kDa and 70 kDa. These bands were not detected in the eluate from Mic10-1-V5 immunoprecipitation after ATPTb2 was targeted by RNAi (Fig. 2B). The former band corresponds to uncrosslinked ATPTb2 and the latter most likely to the aforementioned putative tripartite ATPTb2/Mic10-1/unknown partner adduct. Unlike Mic10-1, C-terminal V5-epitope-tagged Mic10-2 did not co-immunoprecipitated ATPTb2 (Fig. S2A), eliminating this paralog as an interaction partner with ATP synthase.

To investigate if Mic10-1 associates with the entire ATP synthase, rather than with its subcomplex or unassembled ATPTb2, proteins in the elution from co-IP with Mic10-1-V5 were trypsinized and identified by liquid chromatography-tandem mass spectroscopy (LC-MS/MS). Mock IPs on cell lines lacking the V5 tag were performed in parallel as a negative control. The protein enrichment in comparison to mock IP controls was quantified using the label free quantification (LFQ) by a previously described pipeline (38). The identified proteins were filtered to meet the following criteria: 1) presence in all three triplicates; 2) the mean of each triplicate’s Log2-transformed LFQ intensity score >23; 3) absence from at least two out of three mock IPs; 4) presence within the ATOM40 depletome, which indicates that proteins are imported via this outer membrane translocator into the mitochondrion (39). In total, 51 proteins out of 209 detected proteins conformed to these criteria (Fig. 2C-D, S2B, Data set S1). Subunits of the ATP synthase complex and MICOS that are embedded or peripheral to the mitochondria inner membrane (32), were the most represented proteins in the dataset (Fig. 2D). Out of the known 23 subunits that make up the trypanosome ATP synthase, a total of 17 subunits were identified (Fig. 2C), including 5 of 6 subunits of the F_1_. As expected, proteins belonging to the MICOS complex were also identified, with all integral subcomplex proteins being present alongside two components of the peripheral moiety; in total, six out of the nine MICOS proteins fit the criteria set above.

In contrast, only one protein from the electron transport chain complex IV (a.k.a. cytochrome *c* oxidase) was recovered using the same criteria out of the 29 proteins that constitute the complex (40). Two other integral inner membrane proteins found to co-IP with the bait were mitochondrial carrier proteins, one of which was an ADP/ATP carrier protein (41). The other 25 proteins found within the set benchmark consisted mainly of highly abundant enzymes affiliated with the tricarboxylic acid cycle, as well as other prolific soluble mitochondrial matrix proteins (Fig. S2B). To conclude, we present evidence that Mic10-1 interacts with ATP synthase given that the complex’s subunits are enriched in Mic10-1-V5 IPs.

### Trypanosome ATPTb8 is a functional analog of opisthokont F_1_F_O_-ATPase synthase dimer-enriched subunit e

Previous studies in budding yeast demonstrated that Mic10 directly interacts with ATP synthase dimers via subunit e (27, 28), which is essential for the stability of dimers (20, 42, 43) but does not contribute directly to the monomer-monomer interface (44). In Opisthokonts, subunit e contains a conserved GxxxG motif located within its single transmembrane domain (TMD) (43). Analyzing all the ATP synthase subunits identified in the Mic10-1 pulldowns (Fig. 2C), we identified a low molecular weight protein, termed ATPTb8 (45), that contains this motif (Fig. 3A) and is highly conserved among all kinetoplastids (Fig. S3). A search with HHpred (46) revealed a similarity of the region of ATPTb8 encompassing the TMD to subunit e from yeast and human (a.k.a ATP5ME) (Fig. 3A). Additionally, a Kyte-Doolittle hydropathy plot comparison of ATPTb8 and human subunit e shows highly similar hydrophobicity profiles (Fig. 3B). SWISS-MODEL (47) identified mammalian subunit e among the best templates for structure homology modelling. The modelled part of ATPTb8 corresponds to the transmembrane region of the mammalian su-e, including the GxxxG motif, which homotypically interacts with its counterpart on subunit g (Fig. 3C). Just like subunit e, subunit g was implied in dimer stabilization in yeast (48), but recently the protein was also proposed to make interdimer contacts in the rows of mammalian ATP synthases (49, 50).

**Figure 3.**
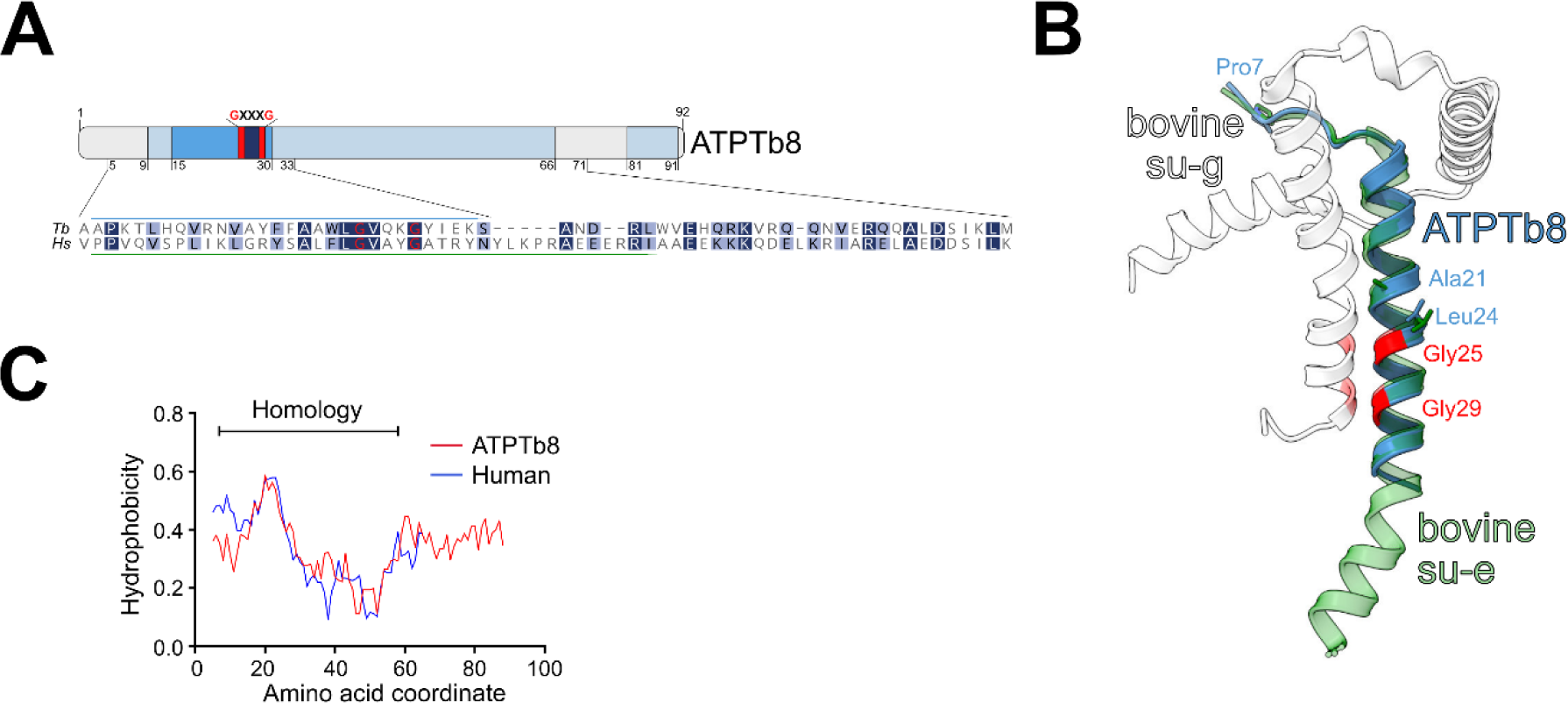
Trypanosome ATPTb8 is a distant homolog or analog of opisthokont ATP synthase subunit e. A) A schematic representation of critical features of the ATPTb8 primary structure. The predicted helical regions and the transmembrane domain are shown in pale and dark blue, respectively. The characteristic embedded GxxxG is labelled. Sequence alignment with human subunit e is shown for the region with similarity revealed by HHPred search. The regions of ATPTb8 and mammalian subunit e depicted in C are marked with blue and green bars, respectively. B) Kyte-Doolittle hydropathy profile comparison of ATPTb8 and human homolog of subunit e. The range of homology between both sequences is shown as the bar above based on HHpred analysis. C) The model of ATPTb8 predicted by SWISS-MODEL using the structure of bovine subunit e (su-e) as a template (PDB ID 6zpo; (44)). The model is superposed with the bovine subunit e and g (su-g) dimer. Glycines of all GxxxG motifs are in red. The conserved residues, which are involved in hydrophobic interactions between subunits e and g, are shown as sticks (44).

Because the bioinformatic predictions indicated that ATPTb8 might be a distant homolog of opisthokont subunit e, we examined whether this subunit is required for the formation or stability of ATP synthase dimers in *T. brucei*. Utilizing 2D protein gel electrophoresis, with denaturing SDS-PAGE following the first dimensional BN-PAGE gel after treatment with 1.5% DDM, we show that C-terminal V5-tagged ATPTb8 is indeed predominantly detected in the dimer fraction (Fig. 4A). Inducible RNAi silencing of ATPTb8 (ATPTb8↓) resulted in cellular growth arrest after 72 hours (Fig. 4B, C). The growth defect in ATPTb8↓ is likely a consequence of compromised ATP production by oxidative phosphorylation and it is consistent with growth phenotypes previously observed after silencing of other trypanosomal ATP synthase subunits (51, 52). No alterations to steady state levels of Mic60 and Mic10-1 were detected in ATPTb8↓ inductions (Fig. S4). Noteworthy, depletion of ATPTb8 preferentially affected stability and/or assembly of ATP synthase dimers, as shown by western blots of BN-PAGE resolved mitochondrial lysates probed with antibodies against subunits β and p18 (Fig. 4D).

**Figure 4.**
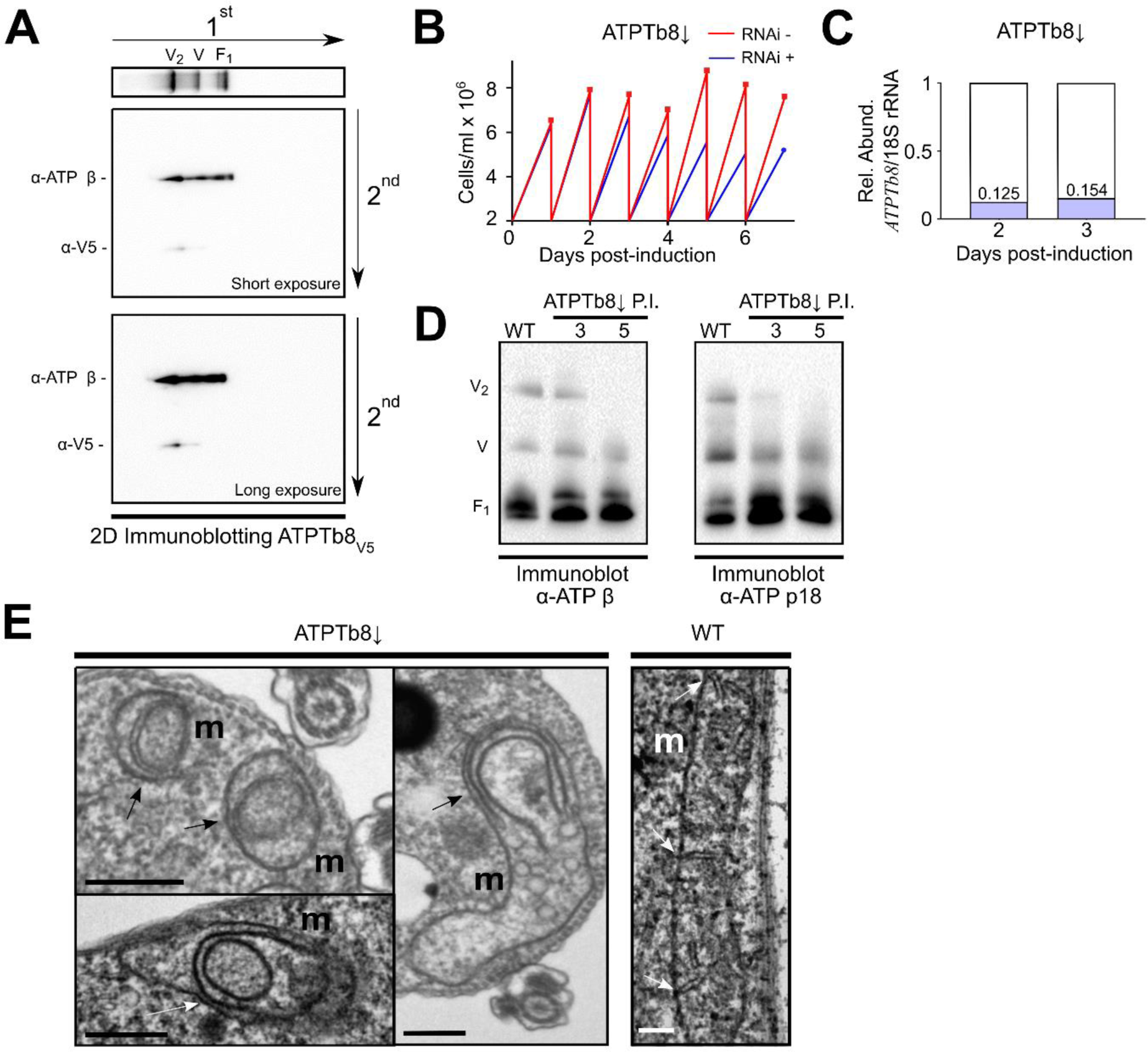
ATPTb8 is enriched in with F_O_F_1_-ATP synthase dimers and crucial for cristae formation. A) Two-dimensional protein gel electrophoresis of ATPTb8-V5, with denaturing SDS- PAGE following the first dimensional BN-PAGE. First dimension immunoblot depicted on top against α-ATP β, showing positions of ATP synthase dimers (V_2_), monomers (V) and free F_1_ moiety. Second dimension immunoblot depicted below against α-ATP β and α-V5 at both short (Top) and long (Bottom) exposures. B) Measurement of ATPTb8↓ and negative control cell growth in glucose-rich medium in which cells are diluted to 2 x 106 cells/mL every day (n = 3; error bars, SD). Cell density, y axis; Days post-induction, x axis. C) Quantitative PCR verification of ATPTb8↓. The relative abundance of *ATPTb8* mRNA in RNAi-induced cell lines in comparison to uninduced control cells were normalized to the unaffected 18S rRNA at 2 and 3 days post-induction. D) BN-PAGE of 1.5% n-Dodecyl-β-D-Maltoside solubilized mitochondria from ATPTb8↓ at 3- and 5-days post induction (P.I.) compared to wild type (WT). Immunoblots against F_1_ subunits α-ATP β and α-ATP p18 shown. E) Transmission electron micrographs (TEM) of WT *T. brucei* and ATPTb8↓ 3 days post induction. Arrows point towards cristae of mitochondria (m). Scale bars, 500 nm.

To investigate whether loss of ATPTb8 has an effect on mitochondrial morphology, transmission electron microscopy of cell sections was performed on cells after 3 days of RNAi induction. Mitochondria exhibiting loss of this subunit depicted anomalous cristae that had circular or semi-circular shapes (Fig. 4E), reminiscent of the onion-like structures that appear upon subunit e deletion in budding yeast (20, 25). Collectively these data suggest that ATPTb8 is a dimer-incorporated subunit much in the same vein as subunit e in budding yeast.

### Trypanosome Mic10-1 interacts with dimeric F_1_F_O_-ATP synthase enriched with ATPTb8

To confirm that Mic10-1 interacts with fully assembled ATP synthase dimers in *T. brucei*, ATPTb8 was *in situ* C-terminal V5-epitope-tagged to perform a crosslink IP. Because ATPTb2 coimmunoprecipitated with ATPTb8-V5, the tag most likely does not interfere with the incorporation of the protein into ATP synthase (Fig. 5A). Next, we probed the IP eluate for the presence of Mic10-1, which was observed mostly as uncrosslinked (Fig. 5B), suggesting that Mic10-1 does not need to be tethered to any interaction partner to co- immunoprecipitate with ATPTb8, unlike with ATPTb2 (Fig. 2A). This further alludes to the possibility that Mic10-1 interacts more firmly to ATP synthase dimers enriched with ATPTb8. Interestingly a faint band at 40 kDa and a stronger band at 70 kDa was also detected, reminiscent of the Mic10-1 immunoband pattern seen in the ATPTb2 IP (Fig. 2A).

**Figure 5.**
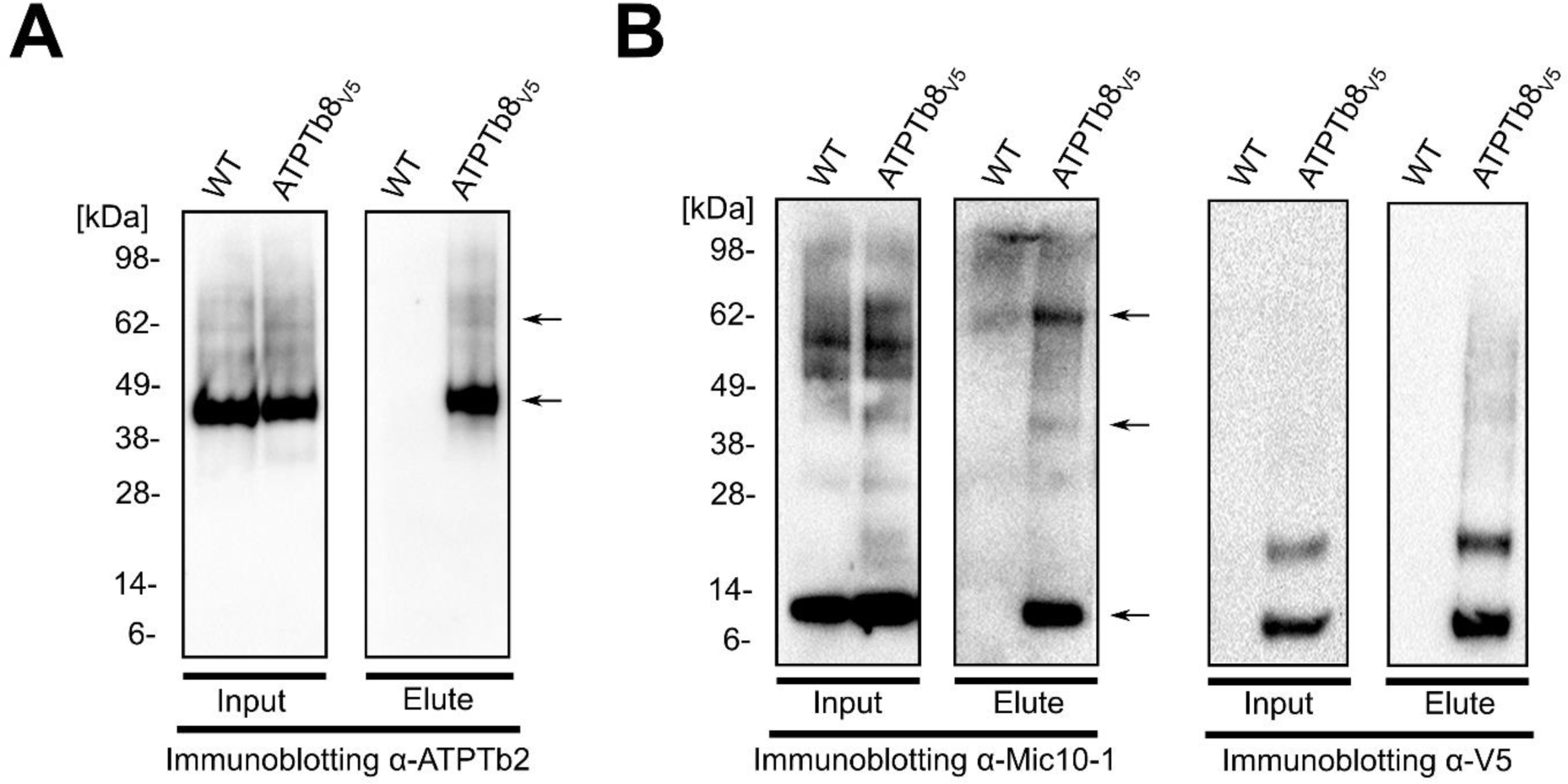
Mic10-1 interacts with F_O_F_1_-ATP synthase dimers. Immunoprecipitation of WT and ATPTb8-V5 mitochondria crosslinked as described in Figure 2. A) Input (10% of elute) and eluates were resolved by SDS-PAGE and immunoblotted using α-ATPTb2 antibody. Arrows indicate discernible bands corresponding to ∼45 kDa (ATPTb2) and ∼70 kDa. B) Same as in A except using antibody against Mic10-1 and V5 peptide. Arrows indicate discernible bands corresponding to ∼10 kDa (Mic10-1), ∼40 kDa, and ∼70 kDa.

## Discussion

Crosstalk between MICOS and ATP synthase was first demonstrated in the budding yeast *Saccharomyces cerevisiae*, a model organism representing the supergroup Opisthokonta, which also encompasses humans. Deletion of Mic60 leads to an accumulation of nonionic- detergent resistant ATP synthase oligomers, likely connected to the apparent functional antagonism between Mic60 and the ATP synthase dimerization subunits e and g in yeast (29). At least a fraction of Mic10 interacts with subunit e, which appears to promote ATP synthase dimer oligomerization (27, 28). Indeed, these two phenomena were proposed to be linked in the Mic60 deletion mutants: the cumulative effect of MICOS disruption with the remaining Mic10’s stabilizing activity on ATP synthase dimer oligomers resulting in the observed hyper-oligomerization phenotype (3, 27).

The physiological role of MICOS-ATP synthase crosstalk in yeast mitochondria remains enigmatic (3), precluding any rational means to hypothesize whether this can be a general eukaryotic phenomenon and not just a fungal novelty. In this work, we have documented the interplay between MICOS and ATP synthase dimers in *T. brucei*, an experimental model belonging to the supergroup Discoba, which separated from Opsithokonta ∼1.8 billion years ago (22, 30). Even though particular aspects of these complexes are conserved between yeast and trypanosomes, such as MICOS maintaining cristae junctions (31) and rows of ATP synthase dimers occurring at the rims of discoidal cristae (33), both complexes exhibit numerous divergent features (29, 37). Based on early branching of the two lineages and marked divergence of their MICOS and ATP synthase, we reason that the crosstalk between MICOS and ATP synthase is a fundamental and ancestral property of cristae.

Specifically, we have demonstrated that Mic60 depletion in *T. brucei* phenocopies the stabilization of ATP synthase oligomerization observed in *S. cerevisiae* Mic60 deletion mutants. In addition to being our first indication that there is MICOS-ATP synthase crosstalk in *T. brucei*, this result also supports the putative designation of this subunit as Mic60 despite its lack of a mitofilin domain (31).

We show that crosstalk between MICOS and ATP synthase is mediated by one of the two trypanosome Mic10 paralogs, Mic10-1. We demonstrate that Mic10-1 crosslinks to ATPTb2 and ATPTb8, two membrane-associated subunits of F_O_ moiety (34, 37). This observation is consistent with Mic10-1 being an integral membrane protein mostly comprised of two TMDs (31). Furthermore, Mic10-1 co-IPs most of ATP synthase subunits after crosslinking. However, ATP synthase subunits were scarcely detected when MICOS subunits were IPed in the absence of any crosslinker (31), suggesting this inter-complex interaction is dynamic and perhaps transient (53).

Our results demonstrate that there are functional differences between Mic10-1 and Mic10-2, as the latter does not interact with ATP synthase. Because Mic10-1 mediates this crosstalk as Mic10 does in yeast, it may represent the conventional Mic10 paralog while Mic10-2 may be the diversified variant whose precise role in discoidal cristae shaping remains undefined.

Indeed, trypanosome Mic10-1 has the typical GxGxGxG glycine-rich motif in the C-terminal TMD, a property shared with other Mic10 homologs, whereas Mic10-2’s corresponding TMD has a reduced GxGxG motif (31). It is tempting to speculate that this difference may be responsible for Mic10-1 interacting with ATP synthase and explain why Mic10-2 does not. However, other factors instead of or in synergy with the GxGxGxG motif could possibly mediate Mic10’s interaction with ATP synthase.

Because yeast Mic10 interacts with ATP synthase dimers, we set out to identify any subunits that may potentially affect dimerization and thus serve as a marker for this higher-order configuration. The F_O_ moiety subunits of *T. brucei* ATP synthase identified by mass spectroscopy have not been fully characterized to date (37). Thus, we screened these subunits for the presence of a single pass TMD with a GxxxG motif, a signature of yeast subunit e (23, 43). The protein ATPTb8 emerged from this screen, and outputs of structural modeling and Hidden Markov searches support homology with human subunit e. ATPTb8 was enriched in dimer fractions. Its ablation phenotype is highly evocative to that of yeast subunit e, in which ATP synthase dimers become sensitive to nonionic detergent treatment along with the correlative emergence of defective cristae (24). Whether discoban ATPTb8 and opisthokont subunit e arose through divergent or convergent evolution remains an open question. Phylogenetic analysis may be precluded by short lengths of these polypeptides and a high degree of divergence, as observed in human and yeast homologs of subunit e, despite belonging to the same eukaryotic supergroup.

Why is this crosstalk between MICOS and F_1_F_O_-ATP synthase present in both Opisthokonts and Excavates and what role does it play? Two different hypotheses have been previously given, and here we propose a third (Fig. 6). These hypotheses may not necessarily be mutually exclusive and perhaps may not apply to all lineages. The first hypothesis postulates that an extra-MICOS fraction of Mic10 interacts with ATP synthase dimers. The local negative curvature induced by Mic10 (15, 16) may relieve membrane tension caused by the positive bending mediated by dimer rows (54), stabilizing the ATP synthase oligomers. In addition, Mic10 binding might enable coordination between MICOS and ATP synthase to temporarily and spatially regulate cristae shape (3, 27). In the second hypothesis, Mic10 acts to bridge the MICOS complex with ATP synthase dimers directly at cristae junctions (28). In this scenario, the ATP synthase dimers would induce positive curvature of tubular necks of cristae. However, it should be noted that cristae junctions in *S. cerevisiae* and other fungi have a predominantly slot-like morphology mainly comprised of flat membranes (9). Our third hypothesis is that we are capturing a transient Mic10-1 interaction between ATP synthase occurring at nascent cristae during the process of ATP synthase dimer-induced invagination of the inner membrane at the future site of cristae junctions. This interaction may involve Mic10 alone or as part of a putative MICOS subassembly present on emerging cristae. Why the occurrence of this crosstalk has been preserved throughout the diversification of Eukaryotes is still waiting to be answered.

**Figure 6.**
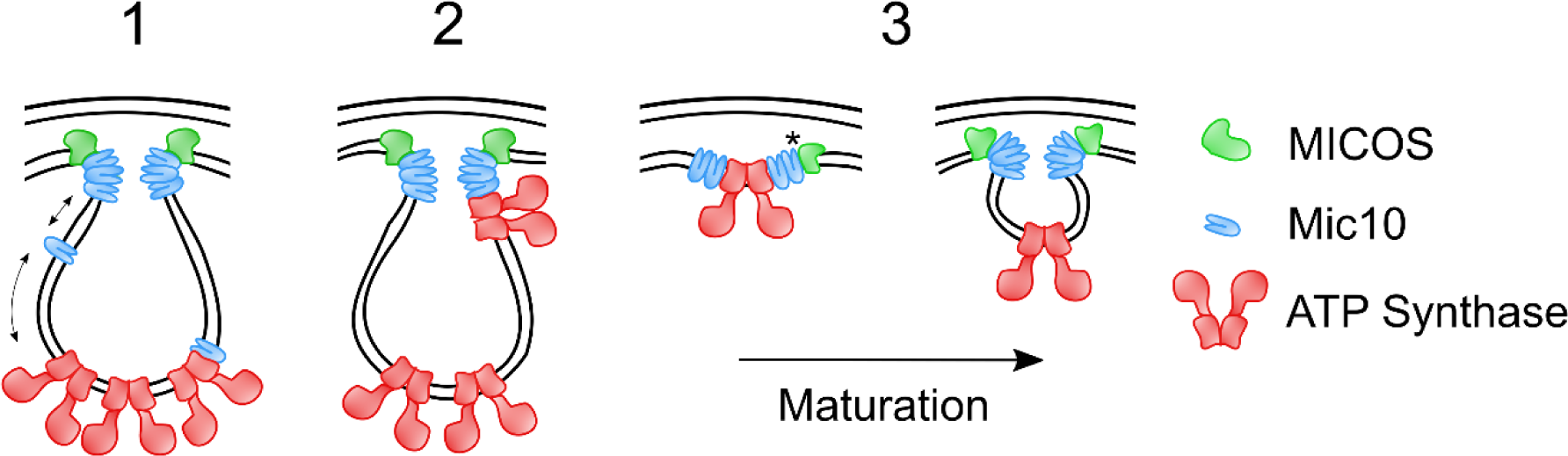
Possible modes for the functional interplay between MICOS and F_O_F_1_-ATP synthase dimers in Eukaryotes. Schematic representation of 3 possible mechanisms of functional interplay between MICOS (green) and F_O_F_1_-ATP synthase (red) in the mitochondrial inner membrane: 1) An extra-MICOS fraction of Mic10 (blue) interacts with ATP synthase dimers at cristae rims. Arrows represent crosstalk between MICOS and ATP synthase dimers via Mic10. 2) Mic10 bridges the MICOS complex to ATP synthase dimers directly at cristae junctions, enabling dimers to induce positive curvature at the tubular necks of cristae. 3) Transient Mic10-1 interacts with ATP synthase at nascent cristae during ATP synthase dimer-induced invaginations. Asterisk represents a putative MICOS complex subassembly. See discussion for more in-depth description of each scenario.

## Material and Methods

### Generation of transgenic cell lines

All trypanosome cell lines used in this study were derived from *T. brucei* SmOxP927 procyclic form cells, *i.e.* TREU 927/4 expressing T7 RNA polymerase and tetracycline repressor to allow inducible expression (55) and are listed in Table S2. The C-terminally tagged ATPTb8-3xV5 and ATPTb2-3xV5 cell lines were generated using the pPOTv4 vector containing a hygromycin resistance cassette by PCR amplification with the oligonucleotides based on an already established protocol (56) in Table S2. The sequence underlined anneals to the pPOT vector.

ATPTb8 RNAi constructs were made by PCR amplifying a fragment of the ATPTb8 gene from *T. brucei* strain 927 genomic DNA with the oligonucleotides listed on Table S2. Utilizing the underlined Xhol and BamHI restriction sites within the primer, these fragments were cloned into the pAZ055 stem-loop RNAi vector (Table S2).

### Trypanosome cell culture

Cells were grown at 27°C in SDM-79 medium supplemented with 10% (v/v) fetal bovine serum and 7.5 mg/L hemin. Cells were grown in the presence and absence of the 1 μg/ml doxycycline for RNAi-induction. Cell density was measured using the Beckman Coulter Z2 Cell and Particle Counter and maintained at exponential mid-log growth phase throughout the analyses.

### SDS-PAGE and Western botting

Protein samples were separated on a Bolt 4-12% Bis-Tris Plus gel (Invitrogen), blotted onto a PVDF membrane (Amersham), blocked in 5% milk and probed with the appropriate primary antibody listed in Table S2 diluted in 5% milk in PBS-T. This was followed by incubation with a secondary HRP-conjugated anti-rabbit or anti-mouse antibody (1:2000, BioRad), depending on the origin of the primary antibody. Proteins were visualized using the Pierce ECL system (Genetica/Biorad) on a ChemiDoc instrument (Biorad).

### Blue Native PAGE, In-gel histochemical staining of F_1_-ATPase activity and 2-D electrophoresis

Blue native PAGE (BN-PAGE) of mitochondrial lysates was adapted from published protocols (34). Briefly, mitochondrial vesicles from ∼5×10^8^ cells were resuspended in 100 μL NativePAGE sample buffer (Invitrogen), lysed with 1.5% dodecylmaltoside (DDM; v/v) for 20 min on ice then cleared by centrifugation (18,600 X g, 15 min, 4°C). The protein concentration of each lysate was determined by Bradford assay, so that 50 μg of total protein could be mixed with 5% Coomassie brilliant blue G-250 before loading on a 3-12% Bis-Tris BNE gel (Invitrogen). After electrophoresis (2.5 hr, 150 V, 4°C), the gel was either blotted onto a PVDF membrane (Amersham) or incubated overnight in an ATPase reaction buffer (35 mM Tris pH 8.0, 270 mM glycine, 19 mM MgSO_4_, 0.5% Pb(NO_3_)_2_, 15 mM ATP). For 2D electrophoresis, individual BNE lanes were cut and placed horizontally on a 12% polyacrylamide gel prior to electrophoresis and western blot analysis.

### Hypotonic mitochondria isolation

Mitochondrial vesicles were obtained by hypotonic lysis as described in previously (31). Briefly, cell pellets from 5×10^8^ cells were washed in SBG (150 mM NaCl, 20 mM glucose, 1.6 mM NaHPO4), resuspended in DTE (1 mM Tris, 1 mM EDTA, pH 8.0) and disrupted through a 25G needle. To reintroduce a physiologically isotonic environment, disrupted cells were immediately added to 60% sucrose. Samples were centrifuged (12,300 X g, 15 min, 4°C) to clear the soluble cytoplasmic material from pelted mitochondrial vesicles. The resulting pellets were resuspended in STM (250 mM sucrose, 20 mM Tris pH 8.0, 2 mM MgCl_2_) and incubated with 5 μg/ml DNase I for 1 hr on ice. An equal volume of STE buffer (250 mM sucrose, 20 mM Tris pH 8.0, 2 mM EDTA pH 8.0) was subsequently added and then centrifuged (18,600 X g, 15 min, 4°C). Pellets enriched with mitochondrial vesicles were then snap frozen in liquid nitrogen for further analysis or subsequently treated with chemical crosslinker, as described below.

### Chemical crosslinking

Hypotonically isolated mitochondrial lysates were resuspended in 1 ml PBS pH 7.5 and incubated with 80 μM DSP for 2 hr on ice. The reaction was stopped by the addition of 20 mM Tris-HCl pH 7.7 for 15 min at RT then cleared by centrifugation (18,600 X g, 15 min, 4°C). The crosslinked mitochondrial vesicles were then snap frozen for subsequent immunoprecipitation (IP).

### Immunoprecipitations

IP of tagged proteins was adapted from published protocols (31). In brief, DSP-crosslinked mitochondrial vesicles from ∼5×10^8^ cells were solubilized in IPP50 (50 mM KCl, 20 mM Tris-HCl pH 7.7, 3 mM MgCl_2_, 10% glycerol, 1 mM phenylmethanesulfonyl fluoride (PMSF), complete EDTA free protease inhibitor cocktail (Rosche)) supplemented with 1% Igepal (v/v) for 20 min on ice. After centrifugation (18,600 X g, 15 min, 4°C) the supernatant was added to 1.5 mg of anti-V5 conjugated magnetic beads, previously washed thrice in 200 μl of IPP50 + 1% Igepal for 5 min at RT. The solubilized mitochondria were rotated with beads for 90 min at 4°C. After removal of the flow through, the beads were washed three times in IPP50 + 1% Igepal. Prior to eluting, the beads were transferred into a new tube. Elution was done in 0.1 M glycine pH 2.0 for 10 min at 70°C and shaking at 1000 rpm. The eluate was neutralized with 1 M Tris pH 8.0. The elution step was repeated to achieve higher recovery. The elutes were further processed for LC-MS^2^ analysis or resolved by SDS-PAGE. IPs were performed in triplicate.

### Protein preparation and Mass Spectroscopy

Triplicate eluates of co-IP proteins were processed for mass spectroscopy analysis as described elsewhere (57, 58). In brief, samples were resuspended in 100 mM TEAB containing 2% SDC. Cysteines were reduced with a final concentration of 10 mM TCEP and subsequently cleaved with 1 μg trypsin overnight at 37°C. After digestion, 1% trifluoroacetic acid (TFA) was added to wash twice and eluates were resuspended in 20 μl TFA per 100 μg of protein. A nano reversed-phased column (EASY-Spray column, 50 cm x 75 μm inner diameter, PepMap C18, 2 μm particles, 100 Å pore size) was used for LC/MS analysis. Mobile phase buffer A consisted of water and 0.1% formic acid. Mobile phase B consisted of acetonitrile and 0.1% formic acid. Samples were loaded onto the trap column (Acclaim PepMap300, C18, 5 μm, 300 Å Wide Pore, 300 μm x 5 mm) at a flow rate of 15 μl/min. The loading buffer consisted of water, 2% acetonitrile, and 0.1% TFA. Peptides were eluted using a Mobile phase B gradient from 2% to 40% over 60 min at a flow rate of 300 nl/min. The peptide cations eluted were converted to gas-phase ions via electrospray ionization and analyzed on a Thermo Orbitrap Fusion (Q-OT- qIT, Thermo Fisher). Full MS spectra were acquired in the Orbitrap with a mass range of 350-1,400 *m/z*, at a resolution of 120,000 at 200 *m/z* and with a maximum injection time of 50 ms. Tandem MS was performed by isolation at 1,5 Th with the quadrupole, HCD fragmentation with normalized collision energy of 30, and rapid scan MS analysis in the ion trap. The MS/MS ion count target was set to 10^4^ and the max infection time at 35 ms. Only those precursors with a charge state of 2-6 were sampled. The dynamic exclusion duration was set to 45 s with a 10 ppm tolerance around the selected precursor and its isotopes. Monoisotopic precursor selection was on with a top speed mode of 2 s cycles.

### Analysis of Mass Spectrometry peptides

Label-free quantification of the data were analyzed using the MaxQuant software (version 1.6.2.1) (59). The false discovery rates for peptides and for proteins was set to 1% with a specified minimum peptide length of seven amino acids. The Andromeda search engine was used for the MS/MS spectra against the *Trypanosoma brucei* database (downloaded from Uniprot, November 2018, containing 8,306 entries). Enzyme specificity was set to C-terminal Arg and Lys, alongside for cleavage at proline bonds with a maximum of two missed cleavages. Dithiomethylation of cysteine was selected as a fixed modification with N-terminal protein acetylation and methionine oxidation as variable modifications. The ‘match between runs’ feature in MaxQuant was used to transfer identification to other LC-MS/MS runs based on mass and retention time with a maximum deviation of 0.7 min. Quantifications were performed using a label-free algorithm as described recently (59). Data analysis was performed using Perseus software (version 1.6.1.3). Only proteins identified exclusively alongside a mean Log2 transformed LFQ intensity scored of >23 and found in the ATOM40 depletome (indicating proteins are imported into the mitochondria) were considered as putative interaction proteins (39). Exclusive identification is here defined as a situation where a given protein was measured in all three replicates of bait protein pulldowns but absent in at least two out of three control replicates.

### Transmission electron microscopy

For ultrastructual studies, cells were centrifuged at 620 X g for 10 min at RT and immediately fixed with 2.5% glutaraldehyde in 0.1 M phosphate buffer pH 7.2. Samples were then post-fixed in osmium tetroxide for 2 hr at 4°C, washed, dehydrated through an acetone series and embedded in resin (Polybed 812; Polysciences, Inc.). A series of ultrathin sections were cut using a Leica UCT ultramicrotome (Leica Microsystems), counterstained with uranyl acetate and lead citrate. Samples were observed using the JEOL 1010 transmission electron microscope (TEM) operating at an accelerating 80 kV and equipped with a MegaView III CCD camera (SIS).

### Bioinformatic analysis

The multiple sequence alignment of ATPTb8 homologs from the order Kinetoplastida was performed by ClustalW using default settings. These alignments were trimmed to remove gaps and regions of poor conservation and rendered in Jalview (version 2.11.1.3) (60). Sequences were obtained from the TriTrypDB database. The homologous regions to human and yeast SuE were determined using HHpred toolkit (46) and the helical region and transmembrane domain within the ATPTb8 sequence were predicted by PSIPRED 4.0 and MEMSAT-SVM software (61), respectively. Kyte-Doolittle hydropathy plots for ATPTb8 and the human su-e sequence were calculated using the ProtScale prediction software courtesy of ExPASy server (62). The structure of ATPTb8 was homology-modelled using SWISS-MODEL (47).

## Data availability

The mass spectroscopy data pertaining to *T. brucei* strain 927 IPs have been deposited to the ProteomeXchange Consortium via the PRIDE partner repository with the dataset identifier PXD025109 (Reviewer access- Username; Password: 3zpMxEB9).

## Acknowledgments

We thank Julius Lukeš for his support in realization of this project, Karel Harant and Pavel Talacko (Laboratory of Mass Spectrometry, Biocev, Charles University, Faculty of Science) for performing LCMS analysis and David Hollaus for technical assistance. Funding from the following sources is gratefully acknowledged: Czech Science Foundation grants 20-23513S to HH, 18-17529S to AZ, and 20-01450Y to OG; Czech Ministry of Education OPVVV16_019/0000759, and Czech BioImaging grant LM2015062.

---

**Figure S1. Depletion of other MICOS subunits does not affect F_O_F_1_-ATP synthase oligomerization. Related to Figure 1.**

A) Immunoblot of *ΔMic10-1*:Mic10-2↓ with antibodies indicated on right. Wild type (WT) and days post induction shown above panel.

B) BN-PAGE of 1.5% n-Dodecyl-β-D-Maltoside solubilized mitochondria from Mic20↓ and Mic32↓ at 5 days post-induction compared to WT. ATP hydrolysis in-gel activity staining shown on the left and immunoblot against α-ATP β on the right. V_2_, dimer; V, monomer; F_1_, free F_1_ moiety.

**Figure S2. Immunoprecipitation of the paralogs Mic10-1 and Mic10-2. Related to Figure 2**.

A) Immunoprecipitation of WT and Mic10-2-V5 mitochondria crosslinked with 80 μM dithiobis(succinimidyl propionate). Input (10% of elute) and eluates were resolved by SDS-PAGE and immunoblotted against α-ATPTb2 and α-V5.

B) Remaining summary of proteins that co-IP with Mic10-1-V5 pulldowns identified by mass spectroscopy within the established threshold and criteria explained in the main text. In green are mitochondrial protein carriers, proteins in grey are components of the tricarboxylic acid cycle (TCA), and in light blue are proteins that cannot be categorized into a single group (n = 3; error bars, SD). Intensity score, y axis.

**Figure S3. ATPTb8 is conserved among kinetoplastids. Related to Figure 3.**

The multiple sequence alignment of ATPTb8 homologs from the order Kinetoplastida was performed with ClustalW using sequences obtained from TritrypDB.org. The GxxxG (GxxxG) motif region indicated by top brackets. Analysis of multiple sequence alignment (AMSA) conservation score and consensus sequence give on the bottom. The following species (accession number, name) were used to generate the alignment: *Crithidia fasciculata* (CFAC1_220013400, ATPCf8), *Leptomonas pyrrhocoris* (LpyrH10_10_0680, ATPLp8), *L. seymouri* (Lsey_0093_0090, ATPLs8), *Endotrypanum monterogeii* (EMOLV88_250011100, ATPEm8), *L. braziliensis* (LBRM2903_250012800, ATPLbr8), *L. panamensis* (LPAL13_250010800, ATPLpa8), *L. enriettii* (LENLEM3045_250011100, ATPLen8), *L. amazonensis* (LAMA_000484200, ATPLam8), *L. Mexicana* (LmxM.25.0590, ATPLmx8), *L. infantum* (LINF_250011100, ATPLin8), *L. donovani* strain LV9 (LdBPK.25.2.000600, ATPLd8), *L. aethiopica* (LAEL147_000405700, ATPLae8), *L. major* (LMJLV39_250011600, ATPLm8), *L. tropica* (LTRL590_250011700, ATPLt8), *L. arabica* (LARLEM1108_250011300, ATPLa8), *L. turanica* (LTULEM423_250011500, ATPLtu8), *L. gerbilli* (LGELEM452_250011200, ATPLge8), *Blechomonas ayalai* (Baya_001_1510, ATPBay8), *Trypanosoma brucei* TREU927 (Tb927.11.600, ATPTb8), *T. congolense* (TcIL3000.A.H_000869900, ATPTco8), *T. vivax* (TvY486_1100440, ATPTv8), *T. rangeli* (TRSC58_00292, ATPTra8), *T. cruzi* (C3747_9g1306c, ATPTc8), *Paratrypanosoma confusum* (PCON_0063970, ATPPcon8), *Bodo saltans* (BSAL_59565, ATPBs8).

**Figure S4. Characterization ATPTb8 RNAi cell line.** Related to Figure 4.

Immunoblot of ATPTb8↓ over time course of RNAi with antibodies indicated on right. Days post-induction (Days P.I.) shown above panel.

**Table S1.** Encoding gene accession numbers and molecular weights of proteins mentioned in this study.

**Table S2.** Key resources table.

**Dataset S1.** List of proteins co-purifying with Mic10-1 after chemical crosslinking. The Perseus software output Excel file of the label free quantification (LFQ) of proteins enriched in Mic10-1-V5 immunoprecipitations (IPs) compared to a mock IP control. This file is appended with: protein names (column A); presence in ATOM40 depletome (column C), signifying being part of the mitoproteome (39); whether a protein fits our criteria for *bona fide* enrichment as described in the main text (column D). Results are filtered for ’Yes’ in column D but can be changed by the user by selecting the appropriate filter criteria accessed in each cell on the top row. Comparison of LFQ scores between Mic10-1-V5 IP (v) and mock IP (c) given in columns E-G. LFQ scores for Mic10-1-V5 and mock IPs for each detected protein given in light blue columns H-N whereas peptide counts are given in grey columns O-Y.The Mic10-1-V5 LFQ scores given in column K (highlighted in green cells with red bold font) were used for bar graphs in Figs. 2C and S2B and summarized in Fig. 2D. Related to Figs. 2 and S2.

